# Divergent Epigenetic and Transcriptomic Reprogramming of Monocyte Subpopulations in Systemic Lupus Erythematosus

**DOI:** 10.1101/2023.12.07.570725

**Authors:** Anna Guiomar Ferreté-Bonastre, Mónica Martínez-Gallo, Octavio Morante-Palacios, Celia Lourdes Calvillo, Josep Calafell-Segura, Javier Rodríguez-Ubreva, Manel Esteller, Josefina Cortés-Hernández, Esteban Ballestar

## Abstract

Systemic Lupus Erythematosus (SLE) is an autoimmune disease characterized by systemic inflammation involving various immune cell types. Monocytes, pivotal in promoting and regulating inflammation in SLE, differentiate from classical monocytes into intermediate monocytes and non-classical monocytes, assuming diverse roles. In this study, we investigated the epigenetic and transcriptomic profiles of these three monocyte subsets in an SLE cohort. In addition to common DNA methylation and transcriptomic alterations, we identified monocyte subset-specific alterations, especially in DNA methylation, which reflect an impact of SLE on the monocyte differentiation process. SLE classical monocytes exhibited a stronger proinflammatory profile, with an interferon signature and were primed for macrophage differentiation. SLE non-classical monocytes displayed a phenotype related to T cell differentiation regulation, and a Th17-promoting phenotype. Changes in monocyte proportions, DNA methylation and expression occurred in relation to disease activity and involved the STAT1 pathway. Integrating bulk datasets with single-cell RNA-seq data of SLE patients further supported the interferon signature in classical monocytes, associating intermediate and non-classical populations with exacerbated complement activation pathways. Our results indicate a subversion of the epigenome and transcriptome in monocyte differentiation toward non-classical subsets in SLE, impacting function, in relation to disease activity and progression.

## Introduction

Autoimmune diseases are chronic inflammatory conditions generally characterized by the presence of autoantibodies and autoreactive T cells. Most research efforts in autoimmune diseases have focused on B and T cells, as the adaptive immune system is pathologically dysregulated. However, inflammation also involves monocytes, macrophages and dendritic cells, which have recently become the focus of many autoimmune disease studies (reviewed in [1]). These cells play a crucial role in the innate immune system, being primarily responsible for phagocytosis of pathogens, antigen presentation and production of cytokines [2]. Monocytes and their derived cells are altered in a wide range of autoimmune diseases, as shown either by their aberrant infiltration to tissues [3,4], causing hyperactivation of adaptive immune system components [5,6], by their presentation of autoantigens to T and B cells [7] or by their defective phagocytosis capacity [8,9].

For years, monocytes were studied as a homogeneous population of immune cells. However, the development of single-cell level techniques, including flow-cytometry, revealed the wide heterogeneity existing within monocytes. The most common classification of monocyte subpopulations is based on the presence of the surface protein CD14, which serves as co-receptor for Toll-Like Receptor 4 (TLR4) and mediates lipopolysaccharide (LPS) signaling, and CD16, also known as Fc gamma receptor IIIa [10,11]. Based on the expression of these two surface proteins, human monocytes were classified in three groups including classical monocytes (cMOs, CD14+CD16-), intermediate monocytes (iMOs, CD14+CD16+) and non-classical monocytes (ncMOs, CD14^dim^CD16+). Indeed, an *in vivo* study by Patel et al [12] convincingly demonstrated that monocyte subsets represent distinct stages of differentiation within the same cell. According to their model, cMOs are released from the bone marrow into the bloodstream, where they reside for about one day before either extravasating to the tissues, undergoing cell death, or differentiating into iMOs. The iMO subset can be found in the bloodstream for up to four days, during which most of them further differentiate into ncMOs, persisting in the bloodstream for up to seven days. The relevance of monocyte subsets has been highlighted by the reported changes in their proportions in many inflammatory conditions such as multiple sclerosis [13], atherosclerosis [14] and, more recently, in COVID-19 [15,16]. This has also been described in SLE, although different studies report opposing results [17–22]

Apart from their differences in cell surface markers, monocyte subsets perform specific functions. In this regard, cMOs are often considered the most inflammatory subset based on their higher production of pro-inflammatory molecules and their propensity to differentiate to dendritic cells and macrophages. iMOs have been reported to perform both pro-inflammatory and anti-inflammatory functions in different contexts. Out of the three subsets, iMOs are the main producers of interleukin-10 (IL-10) upon Toll-Like Receptor (TLR) stimulation [23]. On the other hand, they also are the main monocyte subset that produces pro-inflammatory cytokines in a tumoral environment [24]. Finally, studies of ncMO functions have led to contradicting results: some describe them as the best producers of TNFα upon TLR4 stimulation with LPS [23,25], while others describe them as the least effective to produce pro-inflammatory cytokines [26]. Nevertheless, functions generally associated with ncMOs include maintenance of vascular homeostasis and indicate that they represent the first line of defense against viral pathogens [27–29].

SLE is characterized by multiple organ inflammatory damage and a wide spectrum of autoantibodies. Both genetic predisposition and environmental factors contribute to its development. Genome-wide association studies (GWAS) have revealed many genetic variants conferring susceptibility to SLE, affecting different immune cell types, especially monocytes and other myeloid cells [30]. Particularly, genetic factors seem to have a more definite role in childhood-onset SLE while environmental factors are thought to gain relevance for adult-onset SLE (reviewed in 31 and 32). Environmental factors can trigger changes in the phenotype via epigenetic mechanisms. The most studied epigenetic mark is DNA methylation. In this regard, many studies accumulated in recent years describe aberrant DNA methylation profiles in SLE patients (reviewed in 33). Roughly, these studies report a general loss of DNA methylation in different immune cell types in SLE patients, particularly associated with increased expression of interferon-regulated genes. Type I interferon has a central role in the development and progression of SLE as widely demonstrated by studies showing genetic variance and overexpression of several effectors of these signaling pathways (reviewed in 34). DNA methylation alterations in SLE, and other autoimmune diseases, can both associate with changes in the expression of immune-related genes or serve a sensor of alterations in cellular pathways, becoming useful biomarkers of gene dysregulation [35].

To date, very few epigenetic studies in SLE have focused on monocytes, despite their central role in inflammation. In the present study, we set to inspect the potential existence of monocyte subset-specific epigenetic and transcriptional alterations in SLE, which could shed light on the implication of changes in monocyte subpopulations in the activity and progression of SLE.

## Results

### Monocyte subset-specific DNA methylation profiles in SLE reveal a differential shift in the epigenetic program

We obtained peripheral blood samples from 20 SLE patients when they initially visited the hospital due to a flare episode of the disease or an increase in their symptomatology. Subsequently, we collected new blood samples from these same individuals during their follow-up visits to the doctor. These follow-up visits occurred within a period ranging from 1 to 24 months after the first visit, and 15 of them had achieved remission. Independently, we also collected blood samples from 13 age- and sex-matched healthy donors (HD) (Figure 1A-B). The clinical phenotype data are available in Supplementary Table 1. Blood samples were processed for the isolation of the three monocytes subsets through flow-cytometry cell sorting by following a negative gating strategy similar to previously published studies [13,20,36]. In brief, we first sorted PBMCs negative for CD15, CD3, CD19 and CD56. Subsequently, monocytes subsets were separated based on the surface expression of the markers CD14 and CD16 (Figure 1A).

**Figure 1.**
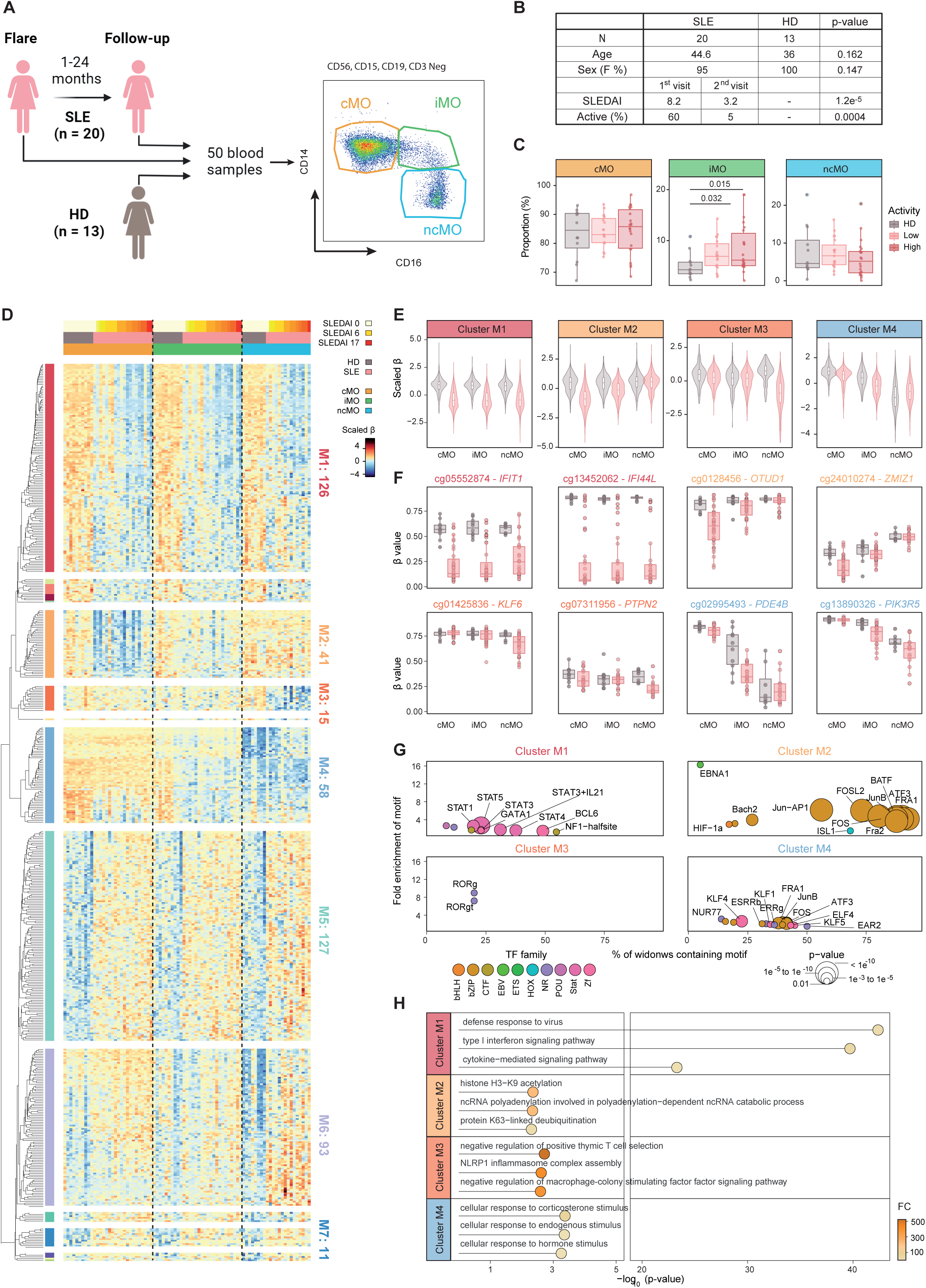
A. Schematic representation of the cohort studied and the obtention of the monocyte subsets. B. Summary table of the cohort characteristics. C. Boxplot representation of the percentage of monocyte subsets within the monocyte fraction of PBMCs. Low activity are samples with SLEDAI of 5 or lower, high activity are samples with SLEDAI of 6 or higher. Statistics performed with two-tailed Wilcoxon test D. Heatmap representation of the 499 differentially methylated positions (DMPs) in either of the pairwise comparisons between SLE and HD samples from each subset. E. Summary violin plot representation of the first four clusters of DMPs. In grey, HD samples, in pink, SLE. F. Illustrative examples of two individual DMPs from the first four clusters. In grey, HD samples, in pink, SLE. The color of the CpG name represents the cluster it belongs to. G. Motif enrichment analysis of the sequence englobing the DMPs. H. Gene Ontology enrichment results of the genes annotated closest to the DMPs from the first four clusters.

The percentages of the obtained monocyte subsets were analyzed, presenting a significant increase in the proportion of iMOs in SLE patients. Notably, this difference appeared to be greater in patients with low activity (SLEDAI < 6) than in patients with high activity (SLEDAI > 6). The proportion of ncMOs also showed differential trends between these two groups of patients, again being relatively higher in patients with low activity (Figure 1C). This was consistent with most of the previous reports, that indicate an increase in CD16+ monocytes. The variability of results associated with disease activity could explain the heterogeneity of previous reports, where some describe an increase in iMOs [18,19] while others reported an increase in ncMOs [20,21].

We then obtained the DNA methylation profiles of cMOs, iMOs and ncMOs of the SLE patient and HD cohorts. The analysis of the results revealed that the different monocyte subsets exhibit unique differences when comparing SLE patients with HDs. Specifically, employing a differential of beta value > 0.1 and an adjusted p-value < 0.2, cMOs displayed 289 differentially methylated positions (DMPs), iMOs exhibited 118 DMPs and ncMOs had 201 DMPs (Figure 1D and Supplementary Table 2). These produced a total of 499 unique DMPs, observing some expected overlap between monocyte subsets. These DMPs were then grouped into seven clusters (M1-M7) through unsupervised clustering based on the DNA methylation levels in the different samples. Consequently, we obtained clusters of DMPs exhibiting a similar behavior across all three subsets and clusters of DMPs displaying a distinctive phenotype in one of the subpopulations (Figure 1E and Supplementary Figure 1A). For instance, cluster M1 consists of 126 DMPs that are hypomethylated in SLE compared to HD across all three monocyte subpopulations. Examples of these DMPs include CpGs annotated to interferon-related genes, such as *IFIT1* and *IFI44L* (Figure 1F). Cluster M2 (41 DMPs) annotate to genes that have a more drastic hypomethylation in cMOs than in iMOs and ncMOs when comparing HD vs SLE (Figure 1D,E). It is the case of genes like *ZMIZ1*, a member of the PIAS (protein inhibitor of activated STAT) protein family, and *OTUD1,* a deubiquitinase related to TNF and IFN signaling [37] (Figure 1F). Cluster M3 (15 DMPs) is particularly interesting as it corresponds to positions hypomethylated in SLE ncMOs in comparison with HD ncMOs (Figure 1D,E).

Examples include those annotating at genes like Krupple-like family transcription factor (TF) *KLF6*, that has been related with the aryl hydrocarbon receptor (AhR) pathway, and *PTPN2* (Figure 1F). Cluster M4 contains 58 DMPs that demethylate in the differentiation from cMOs to ncMOs and that in SLE iMOs are more advanced in the demethylation process than their HD counterparts (Figure 1D,E). Examples of DMPs in the M4 cluster include those annotating to *PDE4B*, encoding a key element for the monocyte’s response to LPS [38] and *PIK3R5*, that encodes a regulatory subunit of PI3K complex, relevant for several immune functions in monocytes including cytokine release and adhesion [39–41] (Figure 1F). The remaining clusters from M5 to M7 (127, 93 and 11 DMPs) were characterized by hypermethylation in SLE in comparison with HD, being the predominant one M5, with similar hypermethylation levels for all monocyte subsets (Supplementary Figure 1A and 1B). In summary, these results suggest that dysregulation of DNA methylation in SLE affects monocyte subsets in different ways. Since iMOs derive from cMOs and ncMOs from iMOs, one can interpret that some determinants in SLE pathology are affecting the differentiation process at the DNA methylation level.

Annotation of the DMPs to the genome in relation to CpG Islands showed different patterns in various clusters (Supplementary Figure 1C). For instance, cluster M1 and M3 showed an enrichment in CpGs annotated to shore regions in comparison to the background. In contrast, the remaining clusters presented a majority representation of positions outside CpG islands. In parallel, annotation of the DMPs in relation to the gene location also separated cluster M1 from the rest (Supplementary Figure 1D). This showed a significant and very marked representation of promoter regions encompassing more than 60% of the DMPs in this cluster. Cluster M3 also showed a high representation of promoter regions as well as exons. The remaining clusters showed a more homogeneous representation with similar region percentages to those present in the background. This highlights the common behavior of the subsets in demethylating promoters while subset-specific changes appear to participate in more complex regulatory processes.

Binding motif enrichment analysis in the regions surrounding DMPs showed that the different clusters present a wide range of associations with different TFs (Figure 1G, Supplementary Figure 1E and Supplementary Table 3). For instance, DMPs from cluster M1 are associated with several STAT TFs possibly indicating that they are related to immune cell activation pathways. Also, around 80% of DMPs from cluster M2 (hypomethylated in cMOs from SLE patients) presented a significant enrichment for TFs of the Fos and Jun family. These have been associated with monocyte-to-macrophage differentiation and activation [42], and they have been previously associated with other inflammatory diseases such as rheumatoid arthritis (reviewed in 43). We were particularly interested in DMPs from cluster M3 because they are hypomethylated in SLE ncMOs, less studied than cMOs. DMPs from cluster M3 presented an enrichment for the consensus sequence of RORγ, which is a TF typically associated with Th17 differentiation [44]. This factor has also been described in a subset of monocytes which is associated with IL-17 production in pathological conditions [45].

We also performed gene ontology analysis of the genes associated by position to these DMPs. The positions showing hypermethylation in SLE in comparison to HD in either cMOs or ncMOs were associated with metabolic synthesis and degradation of several compounds such as “spermidine biosynthesis process” or “allantoin metabolic process” (Supplementary Figure 1F and Supplementary Table 4). Notably, the polyamine spermidine has been previously described to be present at decreased concentrations in the plasma of SLE patients [46]. Also, allantoin is the product of uric acid non-enzymatic oxidation. Urate may be found at elevated levels in the serum of active SLE patients where it facilitates the activation of inflammatory pathways, particularly in those with kidney damage [47]. On the other hand, DMPs that lose methylation in SLE compared to HD in either cMOs or ncMOs were annotated to genes related to immune system activation. DMPs hypomethylated in cMOs were strongly associated with “type-I interferon signaling pathways”, a pro-inflammatory group of cytokines that are a key player in SLE development and pathogenesis. This is consistent with previous results describing strong hypomethylation in interferon pathways in SLE immune cells [48–52].

We also analyzed the gene ontology enrichment of these positions when clustered in an unsupervised manner as in Figure 1D (see Figure 1H, Supplementary Figure 1G and Supplementary Table 5). The results showed that DMPs in cluster M1, which are hypomethylated in SLE in all three monocyte subpopulations, strongly associated with type-I interferon response pathways correlating with our previous analysis. Notably, cluster M3, the cluster particularly hypomethylated in SLE in the ncMO subset, associated with pathways typically associated to T cells and pathways related negatively to monocyte differentiation to macrophages. This pinpointed once more the differential implication of ncMOs in SLE.

### Monocyte subset-specific transcriptomic alterations in SLE

We then performed RNA-seq analysis of 60 samples, corresponding to the 3 monocyte subsets of 13 SLE patients at the first visit and 7 HD. The results showed 2805 differentially expressed genes (DEGs) in cMOs, 1916 in iMOs and 1287 in ncMOs (FDR < 0.5 and a log2FoldChange > 1 or < −1) (Supplementary Table 6). In general, the majority of DEGs corresponded to genes with higher expression levels in SLE than in HDs (Figure 2A-B). Also, as the heatmap representation revealed (Figure 2B), most changes presented a similar behavior in the three monocyte subsets even if a high percentage of the total DEGs did not reach statistical significance in ncMOs. DEGs were divided into seven main clusters (E1-E7) in an unsupervised manner based on their expression levels (Figure 2B-C and Supplementary 2A). In this clustering, DEGs in E1 (986 DEGs) and E2 (631 DEGs) corresponded to genes that show increased expression in SLE versus HDs in all subsets, especially in cMOs. Notably, genes in cluster E2 displayed a progressive decrease in expression during the differentiation from cMOs to ncMOs. In cluster E3 (151 DEGs), genes showed similar expression levels in the three monocyte subsets in HD and a similar upregulation in the three subsets in SLE patients. In cluster E4 (70 DEGs), genes were downregulated during differentiation from cMOs to ncMOs but had a particular increase in expression in ncMOs of SLE. Cluster E5 (971 DEGs) and E6 (375 DEGs) showed a loss of expression in the three subsets in SLE. Finally, cluster E7 (322 DEGs) showed a parallel increase in expression during the differentiation from cMOs to ncMOs in both HD and SLE.

**Figure 2.**
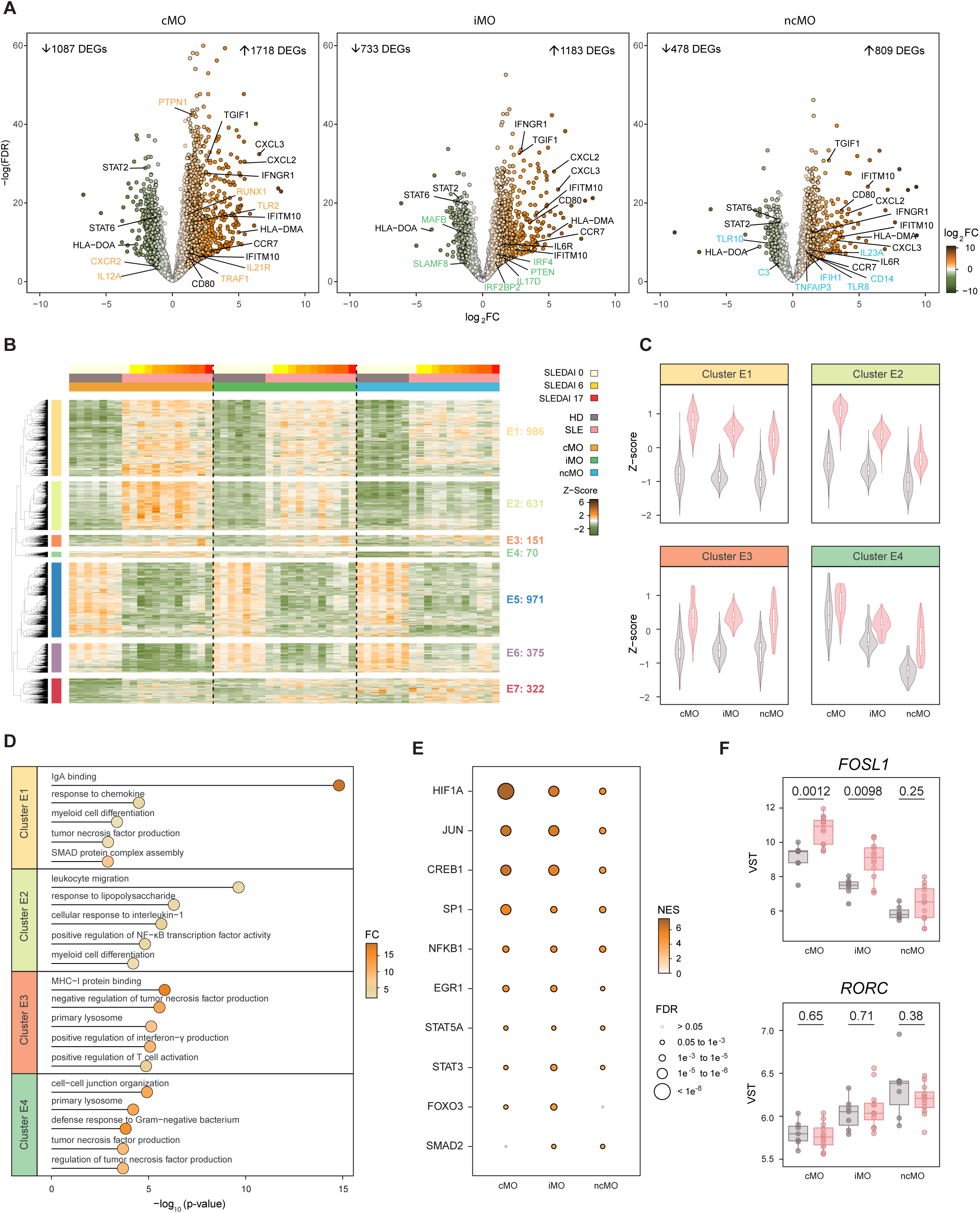
A. Volcano plot of the differentially expressed genes (DEGs) in each pairwise comparison between SLE and HD from each subset of monocytes. Labels in black annotate highlighted DEGs common in the three comparisons. Labels in color annotate highlighted DEGs unique to the respective subset. B. Heatmap representation of the DEGs from the pairwise comparisons between SLE and HD in either of the subsets. C. Summary violin plot representation of the first four clusters of DEGs. In grey, HD samples, in pink, SLE. D. Gene ontology enrichment of the first four clusters of DEGs. E. Dotplot representation results of Virtual Inference of Protein-activity by Enriched Regulon (VIPER) analysis. It displays the top 10 transcription factors predicted to have more activity in SLE samples compared to HD samples, individually in each subset. F. Expression of highlighted transcription factors resulting from the motif enrichment analysis of DMPs. In grey, HD samples, in pink, SLE. Statistics performed with two-tailed Wilcoxon test.

Some genes with a unique behavior in one of the subsets are represented in Supplementary Figure 2B. Among these, *CD58* and *IL18R1* displayed a marked upregulation in cMOs from SLE. CD58 is a molecule expressed in the cell surface of monocytes and it is crucial for the immune synapsis with CD2-expressing T and NK cells [53]. Dysregulation of this axis, through aberrant expression of *CD58*, has been previously linked with several autoimmune diseases including rheumatoid arthritis [54,55]. On the other hand, the proinflammatory cytokine IL18 had increased levels in the serum of SLE patients [56] and the expression of *IL18R* in myeloid cells has been linked with their ability to migrate and be recruited to the site of inflammation [57]. The transcription factor *IRF4*, which we found specifically upregulated in SLE iMOs, is expressed in myeloid cells upon stimulation with IFN-β and it induces their activation and differentiation [58]. *ITGA9*, associated with infiltration and migration in macrophages [59], was found to be especially upregulated in iMOs from SLE patients, indicating a potential migratory phenotype. As for ncMOs, *IFIH1* and *TLR10* were found among the few with a specific differential expression in this subset. Both strongly indicate a pro-inflammatory phenotype in this subset. In the case of *IFIH1*, its expression leads to a strong inflammatory response involving interferons, a known inflammatory pathway dysregulated in SLE [60]. In the case of *TLR10*, its expression in monocytes is linked to a suppression of their activation capacity and their ability to activate T cells [61]. In this case, ncMOs from HD increased their *TLR10* expression in comparison to cMOs, but this increase was not present in the SLE patients which makes them more prone to this interaction than in physiological conditions.

Gene ontology analysis led to the enrichment of biological functions similar to those obtained for DNA methylation (Figure 2D, Supplementary Figure 2C and Supplementary Table 7). Firstly, clusters of DEGs that gain expression (clusters E1-E4) present functions related to immune response. For cluster E1 and E2, particularly upregulated in cMOs, functions related to “leukocyte migration” and “response to chemokine” prevailed. Also, pathways of response through TNF, interleukin-1 and NF-κB were highly represented among the genes from these clusters. Interferon production and MHC-I signaling are predominant among the functions derived from genes from cluster E3. Cluster E4, similarly to cluster E2, showed functions related to TNF production and response to bacteria molecules. For the clusters of genes undergoing downregulation, we observed enrichment in functions related to autophagy (cluster E5), probably related to the deficit of clearance of autophagocytic residues in SLE [8]. In cluster E6, genes related to several metabolic pathways and to the production of IL12, a proinflammatory cytokine produced by myeloid cells downregulated by treatment with corticosteroids [62,63], could lead to a T cell switch towards Th17 population [64].

TF involvement was inferred from the gene expression of their targets using Viper [65]. The TFs with the highest predicted activity in SLE in each of the subsets are shown in Figure 2E. These results were consistent with the observation that the overall transcriptomic profile in SLE is relatively similar among monocyte subsets. HIF1A is the most significantly enriched TF in each of the subsets. This TF has been tightly linked with TNFα signaling and inflammatory autoimmune responses (reviewed in 66). SLE monocytes are in a highly activated state, as also indicated by the enrichment of NF-kB, STATs or EGR1 regulons. NF-kB can be activated through TLR signaling or by uptake of microparticles in SLE becoming a therapeutic target [67–69]. EGR1 has also been shown to associate with inflammatory responses in monocytes [70].

In parallel, we inspected whether the TFs whose binding motifs were enriched in the different DMP clusters (Figure 1G and Supplementary Figure 1C) were also differentially expressed between the populations. This is the case for several of the factors (Figure 2F and Supplementary Figure 2D). For instance, cluster M2 DMPs were clearly enriched for consensus sequences of the family of transcription factors Fos-Jun. FOSL1 is one of the key members of this family and is differentially expressed in cMOs between HD and SLE, but not in ncMOs, showing a similar behavior to the DMPs from this cluster. In the case of cluster M3 of DMPs, where only one TF, RORγ, is significantly enriched in the sequences surrounding the CpG positions, its expression is not differential between HD and SLE in any of the subpopulations. However, it has a clear upregulation between cMOs and ncMOs (Figure 2F), suggesting a more prominent role in the transcriptome regulation of the latter. Other interesting TFs that present differences among the subpopulations include STAT4, ATF3, NR4A1 (*Nurr7*) and IRF4 (Supplementary Figure 2D).

To further study the correlation between the epigenomic profile and transcriptomic profile of the samples, we performed a geneset enrichment analysis of the genes annotated by proximity to the DMPs from each individual subpopulation (Supplementary Figure 2E and 2F). The results showed that the genes annotated to the DMPs from the cMO subset are significantly more enriched in transcriptomic profiles of cMOs from HD samples, in comparison to the SLE samples. Interestingly, cMO’s DMPs were also enriched in the iMOs from HD samples, probably due to the short lifespan of cMOs before they differentiate to iMOs which may cause cMO’s epigenomic profile to influence iMO’s transcriptomics. ncMO’s DMPs did not reach statistical significance threshold for the correlation, but they showed a tendency of being more enriched in the SLE samples.

### DNA methylation changes correlate with SLE activity and progression

We and others have previously shown that the DNA methylation profiles of immune cells correlate with activity index in rheumatoid arthritis [71–73] and in SLE [74]. We set to find whether the various monocyte subsets also met these characteristics. We performed Spearman’s correlation with the SLEDAI of the first-visit samples and their DNA methylation profile. We found 823 CpGs in cMOs, 683 in iMOs and 952 in ncMOs that correlated significantly with disease activity (ρ cutoff = 0.7 & p-value cutoff = 1 × 10e–3) (Figure 3A, Supplementary Figure 3A, B and Supplementary Table 8). These were vastly different between monocyte subsets. However, when analyzing the ontologies of the genes annotated to these positions, the results obtained were similar among the subsets. Particularly, the positions with a negative correlation to SLEDAI, i.e., positions that are less methylated at higher activity indexes than at low activity indexes, associated with pathways related to immune response via type-I interferon pathway in cMOs and iMOs (Figure 3B and Supplementary Table 9). Unfortunately, these positions did not correlate with activity in the second visit samples, in contrast to what we had observed for other autoimmune conditions [71]. This caused very poor correlations when comparing the difference in methylation with the difference in activity (R^2^ < 0.5, data not shown).

**Figure 3.**
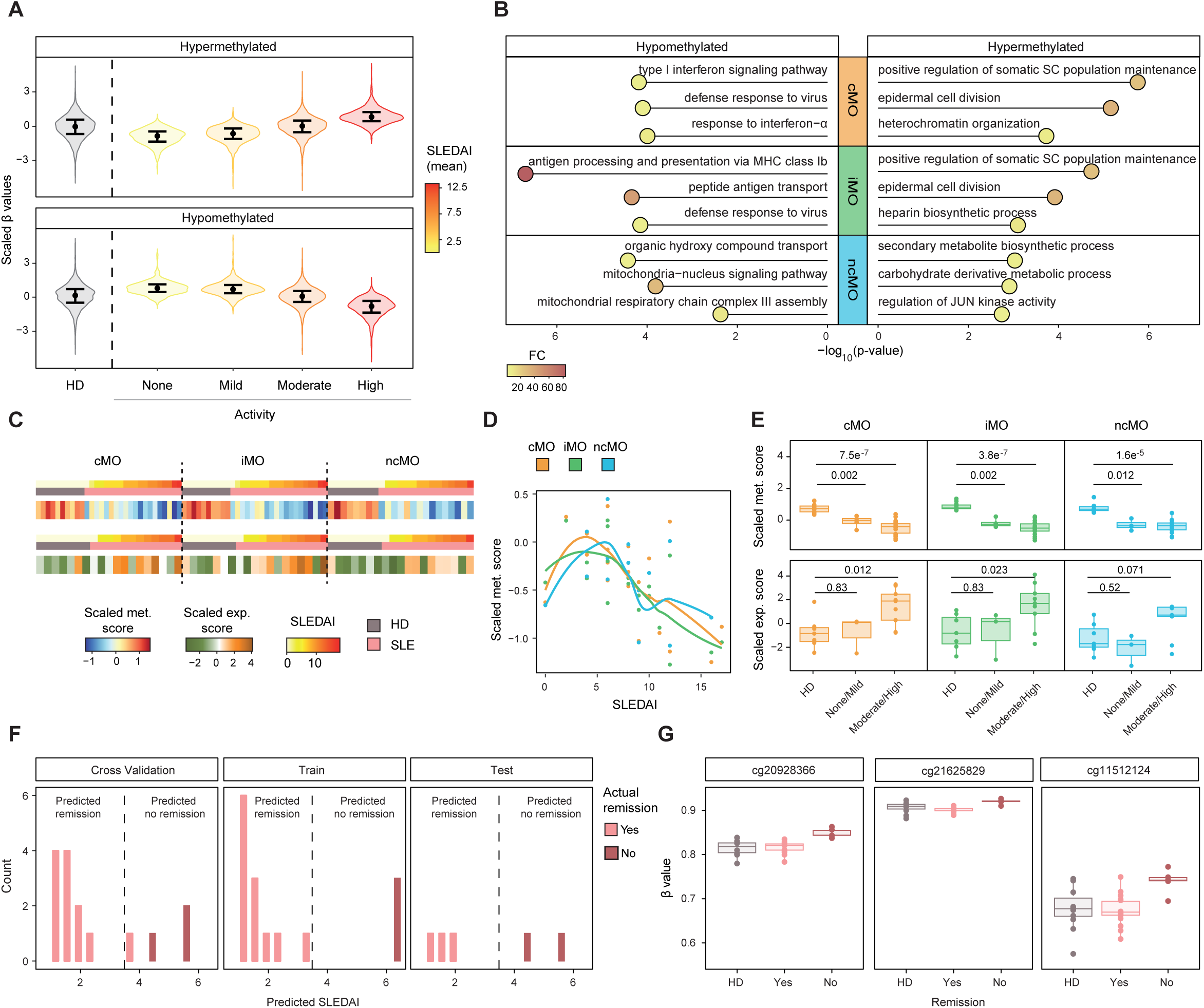
A. Violin plot representation of the cMO’s CpGs correlating with SLE activity as measured by SLEDAI in a Spearman’s correlation. Samples divided in four groups according to their activity index: none is SLEDAI = 0, mild is SLEDAI between 1 and 5, moderate is an index between 6 and 10, and high is a SLEDAI above 10. B. Gene ontology results of the genes annotated by proximity to the CpGs correlated with activity in each of the subsets. C. Heatmap representation of STAT1 targets at the methylation (top) and expression (bottom) level. Methylation targets are defined per HOMER database and expression targets as per CollectRI database. D. Dotplot representation of methylation correlation with SLE activity index of the DMPs that are STAT1 targets in cluster M1. E. Boxplot summary representation of STAT1 targets at the methylation (top) and expression (bottom) level in each of the subsets. Methylation targets are defined per HOMER database and expression targets as per CollectRI database. Samples grouped per activity so that None/Mild group is SLEDAI 0 to 5 and Moderate/High is SLEDAI above 6. Statistics performed with two-tailed Wilcoxon test. F. Barplot representation of the results of the predictive model in the different cohorts. G. DNA methylation levels of the three CpGs used by the predictive model in HD, remission patients and non-remission patients.

Given the ontology results associating the correlating positions with interferon pathways, together with the role of STAT1 in the epigenetic profile (Figure 1G), we wondered whether this individual TF could be associated with the activity in the samples. To address this question, we generated a STAT1 score for both DNA methylation and expression. For DNA methylation, we calculated the average of the scaled methylation values of all the CpGs with the STAT1 consensus sequence in a region of 500bp surrounding the DMPs from cluster M1. For the expression data, we measured the average of the scaled expression values of the genes target of STAT1 as defined by CollecTRI regulons [75]. The results revealed a significant association between the activity index and the levels of STAT1 targets. Samples with higher activity decrease the methylation of STAT1-responsive DMPs while they have a higher expression of STAT1 gene targets (Figure 3C-E). Although the association between the epigenetic STAT1 profile and the activity occurs for the three subsets, the correlation of the transcriptomic profile is only statistically significant in cMOs and iMOs. This suggests that ncMOs as less STAT1-responsive.

We also had information about the progress of the patients. Specifically, 15 out of the 20 patients included in this study improved over the following months and achieved a remission state with a SLEDAI below 6 in the second sample collection point. With this information available, we investigated whether some CpG positions could predict the prognosis of the patients from the first visit DNA methylation status. We identified 494 DMPs in cMOs between patients that would remit and patients that would not. With these, we built a supervised learning k-nearest neighbors’ algorithm to predict the progression of the disease. With an internal 10-fold cross validation approach we confirmed that 3 CpG positions were enough to build an accurate prediction algorithm (Figure 3F and 3G).

### SLE patients undergo drastic changes of monocyte subpopulations

To further investigate the presence of differences in our dataset, we leveraged single-cell RNA-seq dataset from PBMC samples of 162 SLE patients and 98 HDs (76 - GSE174188). Due to the difficulty of identifying iMOs in single-cell datasets, we initially divided the myeloid fraction of this dataset at a very low resolution, obtaining just two clusters: cluster 0 or CD16-cells and cluster 1 or CD16+ cells (Figure 4A-4C). With this initial division, DEGs from our bulk RNA-seq dataset were significantly shared with the DEGs between SLE and HD cells in the single-cell object (cMO’s DEGs in cluster 0, p.value = 0.0096; iMO’s and ncMO’s DEGs in cluster 1, p.value = 0.0164) (Supplementary Figure 4A). Some of the genes upregulated in both datasets in the comparison of HD vs SLE included *IRF7*, *IFI27*, *HLA-A* or *HLA-C* (Figure 4D and Supplementary Figure 4B).

**Figure 4.**
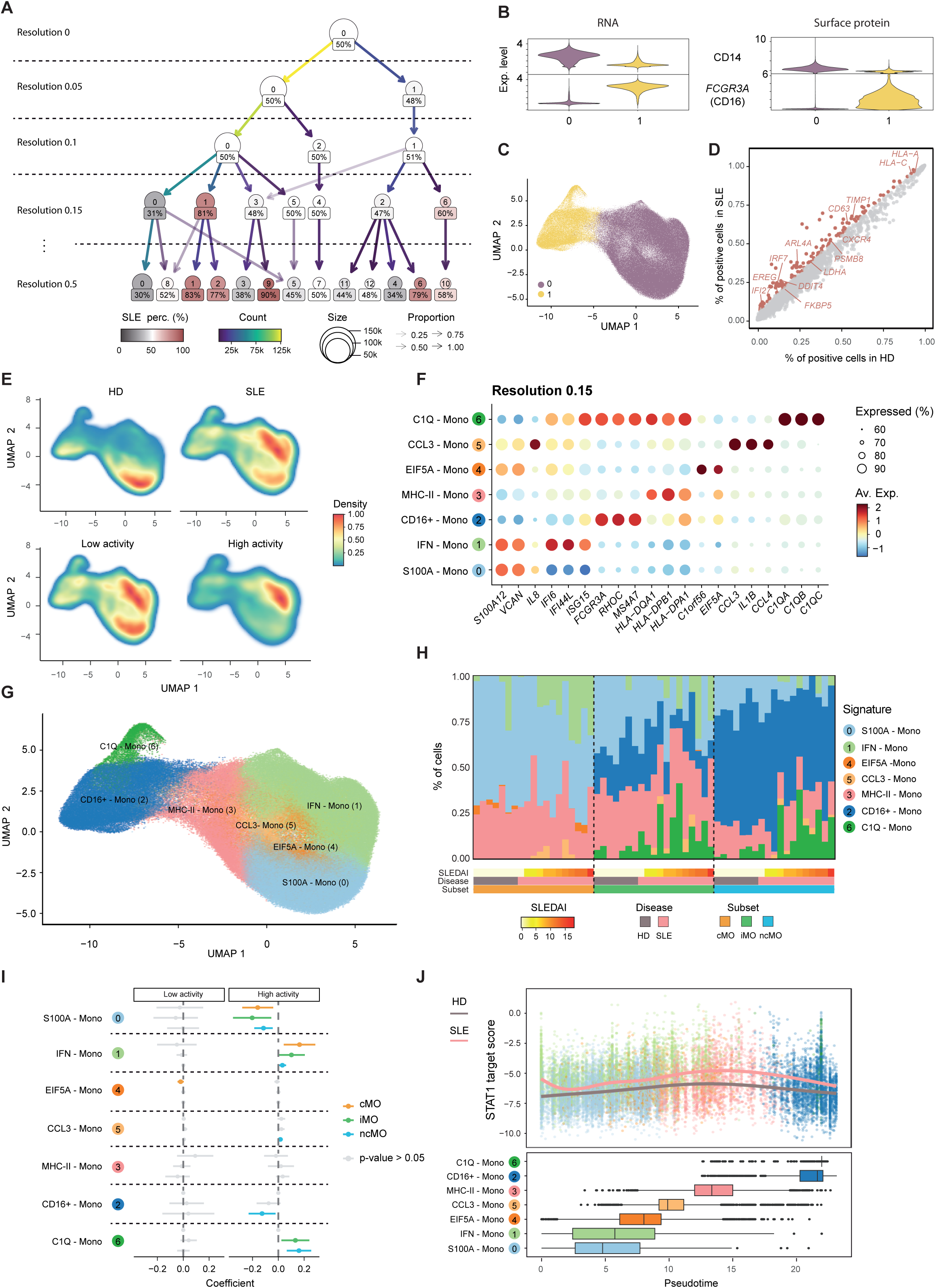
A. Representation of unsupervised clustering results at different resolutions of the myeloid fraction of PBMCs from the single-cell RNA-seq dataset GSE174188. Node color represents the percentage of SLE composition of each cluster. Edge color represents the number of samples following each division. The transparency shows the incoming node proportion. B. Violin plot representation of the expression levels of CD14 and CD16 (*FCGR3A*) at RNA and protein level in the two clusters generated by a 0.05 resolution. C. UMAP representation of the distribution of the two clusters generated by a 0.05 resolution. D. Scatter plot of the percentage of detected genes in HD and SLE samples in Cluster 0 of the resolution 0.05. In color are the DEGs from the sc-RNA-seq dataset (FDR < 0.05, log2FC > 0.1). Labels present in DEGs commonly upregulated in the bulk dataset. E. UMAP representation showing the density distribution of cells in the HD and SLE cohort (top), and within the SLE cohort, the low activity and high activity samples (bottom). SLEDAI of 6 was used as a threshold. F. Dotplot representation of some of the differentially expressed genes in each cluster formed by a 0.15 resolution. G. UMAP representation of the distribution of the six clusters generated by a 0.15 resolution. H. Barplot representation of the estimate percentage composition of each bulk sample according to signature expression from each of the clusters generated by a 0.15 resolution. I. Summary plot of percentage composition of bulk samples. Coefficients, p-values and 95% confidence intervals derived from fitting a linear model and testing it with Wilcoxon test. J. Scatter plot representation of STAT1 targets expression score across the pseudotime distribution of the differentiation from CD16- to CD16+ monocytes (top). Summary representation of pseudotime representation of each cluster (bottom).

Density distribution of cells in the myeloid object was strikingly different between SLE and HD samples. This difference was particularly accentuated in the SLE samples with higher activity (SLEDAI > 6) (Figure 4E). This finding prompted us to divide the object to a higher resolution, looking for clusters particularly composed of one condition of samples (Figure 4F). At a resolution of 0.15 (Figure 4A), two of the new clusters are composed mostly of cells coming from the SLE samples. In particular, clusters 1 and 6 are composed of 81% and 60% SLE cells, respectively (Figure 4A).

Analysis of some of the highly expressed genes for each cluster and their corresponding enriched pathways (Figure 4F, G and Supplementary 4C) led us to the following annotation: Cluster 0, primarily formed by cells of HD origin, was labelled as S100A - Mono and was enriched in pathways of response to lipopolysaccharide but also of response to oxidative stress. Cluster 1, essentially composed of SLE cells received the name of IFN – Mono because it had an unmistakable interferon signature with *IFI6*, *IFI44L* and *ISG15* genes as its top markers. Cluster 2 was the main CD16+ cluster and, therefore, we labelled it as CD16+ - Mono. Cluster 2 was also enriched in pathways of response to type-I interferon and regulation of leukocyte activation. Cluster 3 (MHC-II – Mono) was strongly enriched for genes and pathways of the MHC-II complex and Cluster 4 (EIF5A - Mono) had higher levels of cell proliferation genes (*EFI4A* and *C1orf56*) and pathways related to cell adhesion and endocytosis. Cluster 5 (CCL3 - Mono) had a highly proinflammatory signature with marked TNF-a pathway activation. Finally, Cluster 6 (C1Q – Mono) highly expressed genes and pathways of the complement 1q (C1QA, C1QB, C1QC). Interestingly, MHC-II – Mono, EIF5A – Mono and C1Q – Mono presented very similar features to those found in a previously annotated single-cell dataset of another autoimmune disease [77].

By using CIBERSORTx software [78], we studied the proportion of these clusters represented in our bulk dataset (Figure 4H). With this approach, we determined that our bulk samples are composed of a heterogeneous mixture of cells showing differential patterns correlating with disease state and disease activity. As expected, cMOs from SLE donors were composed by a significant percentage of the labelled IFN - Mono cluster, while HD samples barely harbored them. This representation was more elevated in samples with higher disease activity, where the difference in proportions was statistically significant. We determined that iMOs were made of a mixture of mostly S100A - Mono, MHC-II – Mono and CD16+ - Mono in HD samples while iMO SLE samples repeatedly showed a non-trivial representation of C1Q – Mono. This behavior was similar also for ncMO samples, where C1Q – Mono has a significantly higher proportion in the SLE samples of our bulk dataset. Another remarkable finding is the presence of the highly proinflammatory cluster CCL3- Mono exclusively in SLE samples, being present in all three subsets from the bulk but never in HD donors.

Finally, to inspect the impact of STAT1 dysregulation, initially determined from the bulk methylome and transcriptomic datasets, we studied the STAT1 regulon in the single-cell RNA-seq dataset. As anticipated, we observed an increased pattern of activation of this pathway in SLE samples. This activation was particularly exacerbated in the MHC-II - Mono cluster and less in the S100A – Mono or the CD16+ clusters (CD16+ - Mono and C1Q – Mono) (Figure 4I). Together, these results correlate with our expression results indicating a higher STAT1 influence on the higher activity samples, mainly in the cMO and iMO subsets, which are the samples with a heightened ratio of the MHC-II – Mono cluster.

## Discussion

In this study, we have exploited the power of multi-omics to dissect changes in monocyte subpopulations in SLE. Previous studies had suggested a differential role for CD16+ monocytes through variations in their proportions and functions with respect to those in healthy individuals [17–22]. Our study sheds light into both the differential phenotype of monocyte subsets in SLE and the expansion of certain subpopulations. On the one hand, DNA methylation profiles presented changes between SLE and HD individuals that are common to all three subsets but also display unique differences only present in one of the monocyte subsets. On the other hand, transcriptomic analyses showed that cMOs were the subset with higher and more significant differences between SLE and HD. These differences, both at epigenetic and transcriptomic levels, correlated with disease activity, particularly the STAT1 pathway. We also developed a predictive model that, by making use of the methylation value of only three CpG, could predict the prognosis of the disease. Finally, by applying the information obtained from a public single-cell RNA-seq of a SLE cohort, we discerned the composition of our bulk data and identified a signature of cells that explain the phenotype observed in the SLE monocyte subsets.

Involvement of monocyte subsets in SLE (and other inflammatory conditions) has been subject of debate in the past years. Discrepancy of results in monocyte proportions is patent, where some studies indicate higher percentage of cMOs [17], others an increase in iMOs[18,19] or in ncMOs [20,21] and, finally, other studies have not found differences in the proportions of the subsets between SLE samples and HD [22]. Our results indicate that the proportion of monocyte subsets varies in SLE in relation to disease activity. The proportion of iMOs is increased in SLE samples, particularly in those with low disease activity. The ncMO subset follows a similar behavior, without reaching statistical significance, probably due to cohort size. This suggests a differential role of the subsets at various stages of the disease.

DNA methylation changes can reflect the influence of the environment and impact cellular phenotype, priming cells for a subsequent response to the stimuli received. Continuous systemic inflammatory conditions present in SLE shape the cells of the immune system into a constant state of responsiveness and further inflammation. Multiple studies have shown DNA methylation alterations in SLE patients (reviewed in 33), generally pointing towards a dysregulated type-I interferon signature. These modifications are associated with disease activity and prognosis. Our study on monocytes subsets has allowed us to determine changes in positions shared among the different subsets, indicating a widespread inflammatory behavior dominated by this cytokine. However, we also observed DMPs that are exclusive to each monocyte subpopulation. For hypomethylated DMPs unique to cMOs, up to 80% harbored the consensus sequence for the binding of FOS/JUN family of TFs. In the myeloid compartment, these TFs and the demethylation of their target sequences are associated with differentiation to macrophages [79,80]. This suggests a priming of these cells towards macrophage differentiation, which would take place instead of the differentiation towards iMOs and ncMOs. In contrast, for DMPs exclusively demethylated in SLE ncMOs, we identified an association with T cell regulatory factors and pathways. Some of these DMPs were associated with the TF RORγt, a factor typically found in T cells, which has also been described in monocytes under some pathological conditions [45]. In this report, the expression of RORγt by a subset of monocytes was associated with production of IL17, a cytokine tightly related to the differentiation of Th17 cells [81]. Interestingly, this type of response, particularly fostered by IFN-α conditioned monocytes [82], has been shown to play a role in SLE pathogenesis [83,84]. It is possible to speculate that the observed differences in monocyte subsets could be due to the differential response because of the cells expressing different receptors, of them being primed differently or that the bulk samples are composed of a heterogeneous mixture of different cells.

Transcriptomic analysis showed the predominance of alterations shared by the three subsets, with subset-specific changes mainly displayed by cMOs. This subset presented more and higher differences in gene expression than the rest of subsets. This result probably attests to the fact that cMOs are a less differentiated state of the cells and thus are more plastic and able to respond to an inflammatory environment. As anticipated, the genes upregulated in SLE were related to pro-inflammatory and antigen presentation pathways, whereas the downregulated were associated to negative regulation of autophagy or production of IL12. These are relevant pathways associated with the development of SLE, where the dysregulated autophagosomic system may have an important role the development and severity of the disease (reviewed in 84) and IL12/IL23 axis is important for Th17 cell differentiation [82]. Interferon signature, a well-known driver of SLE, was also present in our samples, particularly in those with higher activity as seen in both epigenetic and transcriptomic profile of STAT1 targets. At the transcriptomic level, it appears that cMOs and iMOs are the most responsive and susceptible subsets, in contrast to ncMOs where STAT1 targets expression does not reach a significant increase in high activity samples.

Leveraging the characteristics of our cohort and the clinical data gathered about progression of disease, we classified our patients into good prognosis or remission and bad prognosis or not remission. With this information, we developed a predictive model that used the epigenetic information from the first visit sample to foresee the prognosis of the samples into these two categories. With the methylation state of just three CpG sites, we were able to achieve this classification in our test and validation cohorts. This suggests that, even at the beginning of a flare period, there are measurable indicators of the prognosis of the episode. A limitation for this part of our study is the size of our cohort, which had to be divided into independent groups for validation and test. Although these results should be further validated with more samples, they are already promising, especially given the sample size.

Finally, using a single-cell RNA-seq dataset from a SLE cohort, we identified subgroups of monocytes within our bulk dataset that help explain the differences observed. In this regard, cMOs and iMOs from SLE samples were formed by a high percentage of cells with an IFN signature, particularly in the high activity samples, which could explain the STAT1 signature that we had identified. Similarly, iMOs and ncMOs also contained a higher percentage of another group of cells with a high C1Q signature. This group of cells is likely to be important in SLE pathogenesis, where immunocomplexes formed by autoantibodies and complement factors are the main initiators of tissue damage. Of note, despite the results show that the bulk SLE samples are formed by a higher percentage of these proinflammatory groups of cells than the HD samples, they also contain an important representation of “healthy” cells or cells present in healthy donors, it is not the whole of monocytes that are pathogenic. Thus, it would be interesting to further study which signals are able to influence a healthy monocyte just released from the bone marrow to develop a pro-inflammatory, disease promoting cell phenotype, or to follow a physiological destiny even in the inflammatory environment present in this disease.

## Methods

### Sample collection

The study was approved by the Committee for Human Subjects of the local ethics committee (PR275/17). 20 Systemic Lupus Erythematosus (SLE) patients and 13 healthy donors were included in the study. Participants gave both oral and written consent for their blood to be used for research purposes. At diagnosis, all SLE patients fulfilled at least 4 EULAR/ACR for the diagnosis of SLE [86]. Samples from each SLE patients were collected at two different timepoints, the first one at the onset of a new flare and the second one at the subsequent visit. For SLE patients, the Systemic Lupus Erythematosus Disease Activity Index (SLEDAI) [87] was registered at each extraction date.

### Sample processing and monocyte isolation

25 ml of whole blood were processed within 24h of collection by laying on Lymphocyte Separation Solution (Rafer, Zaragoza, Spain) and centrifuging without breaking in order to obtain peripheral blood mononuclear cells (PBMCs). Remaining erythrocytes were lysed with ACK lysis buffer. PBMCs were cryopreserved in FBS with 10% DMSO to gather several samples for flow cytometry sorting on the same batch. The day of cytometry sorting, samples were thawed at 37°C and stained for CD19-FITC (BD-Bioscience), CD15-FITC, CD3-FITC, CD56-PE, CD16-APC and CD14-APCVio 770 (Miltenyi). Finally, DAPI was included for the selection of viable cells. Firstly, we filtered for singlets, FSC-SSC myeloid-like cells and DAPI negative cells. Then a negative gating for CD15, CD3, CD19 and CD56 was performed as recommended in [20,36]. Finally, monocyte subsets were separated by their expression of CD14 and CD16 (Figure 1A).

### DNA and RNA extraction and preparation

Sorted samples were pelleted and frozen in RLT Buffer + 10% β-mercaptoethanol as recommended by the kit Allprep DNA/RNA Micro, Mini (Qiagen). After collecting all the samples, double extraction of DNA and RNA was performed in as few batches as possible following manufacturer’s instructions.

For DNA, the three monocyte subsets samples from 20 patients in the first and second visit as well as 10 samples from HD were used, in total 150 samples were analyzed. 250ng of genomic DNA were modified with bisulfite with EZ DNA methylation Gold kit (Zymo Research) following manufacturer’s instructions. Modified material was hybridized in Infinium MethylationEPIC arrays. Since these arrays only fit 8 samples each, a sample distribution strategy was designed to avoid potential batch effects confounded. In this way, disease state, subset, donor’s age and sex, visit number, SLEDAI activity as well as time the sample spent frozen were distributed in a balanced way across arrays [88].

For RNA, the three monocyte subsets samples from 13 patients in the first visit as well as 7 samples from HD were used, in total 60 samples were sequenced in a 100-bp paired-end manner. Libraries and sequencing were performed by BGI Genomics (Hong Kong) with Low input Transcriptome sequencing - Smart Seq based method and DNBseq platform. Approximately 50 million reads were obtained per sample. All samples passed sequencing quality control performed with FastQC [89].

### DNA methylation analysis

Data from DNA methylation studies were analyzed following the pipeline described for the shinyÉpico package [90]. In brief, after removing CpHs, SNPs and X/Y chromosomes, samples were normalized using noob+quantile algorithms and beta values were transformed to M values with lumi package v2.48 [91]. Quality control was performed and sample composition was checked with the function *estimateCellCounts* from minfi package v1.42 [92], some samples were removed due to undesired scores in either of the tests, all from the ncMO subset. In the end, 137 samples were retained for further analysis. Statistical tests were performed with Limma package v3.52 [93] using the arrayWeight argument. Comparisons between SLE samples from the first visit and HD were performed in each subset individually and CpGs with a difference in beta value > 0.1 and an FDR < 0.2 were considered differentially methylated positions (DMPs). Plots were generated with ggplot2 v3.4 and gplots v3.1 packages.

Annotation of CpGs to their closest gene was performed with the function *annotatePeak* from the package ChIPseeker v1.32 [94,95]. Motif Enrichment analysis was performed with HOMER v4.11 [96] with the function *findMotifsGenome.pl,* within a window of 500 around the DMP. For this analysis, the arguments –*cpg* and –*nomotifs* were used and the background was considered as the total of CpGs included in the analysis. Gene Ontology Analyses were performed with rGREAT package [97] with argument *rule* set to twoClosest genes and *version* of GREAT set to 4. Annotation to CpG island context was performed with annotatr package v1.22 [98] and annotationHub v3.4 [99], where shores are defined as 2kb upstream/downstream of CpG Islands, shelves are 2kb further from shores and the rest is outside CGI. Annotation to genetic context was performed with ChIPseeker’s function *annotatePeak*. Correlation of methylation and activity was performed with a Spearman’’s correlation with an estimate threshold of 0.7 and p-value < 0.001.

### Bulk RNA-seq analysis

Files were aligned to hg38 and normalized with Kallisto [100]. Sample SLE15 was excluded from further analysis due to outlier profile. Differential analysis was performed with DESeq2 [101] in each subset individually by comparing HD samples versus SLE samples. A batch effect was detected in some of the HD samples that were sorted and purified separately from the others including HD24, HD6, HD27 and HD28, this was then included as a covariate in posterior analyses. Differentially expressed genes (DEGs) were found by *lfcShrink* function with Ashr algorithm. DEGs were defined as protein-coding genes with an absolute log2FC > 1 and an FDR < 0.05. Variance Stabilizing Transformation (VST) values and normalized counts provided by DESeq2 were used for visualization purposes.

Plots were created with ggplot2 package [102]. Gene Ontology analyses were performed with *enrichGO* function from clusterProfiler package [103] with org.Hs.eg.db database [104]. Virtual Inference Protein-activity by Enriched Regulon Analysis (VIPER - 65) was performed with the dataset from the Collection of Transcriptional Regulatory Interactions [75] with the genes ranked by their p-adjust and log2FoldChange. Gene set Enrichment Analysis (GSEA) was performed with function GSEA from clusterProfiler [103] with the genes ranked by adjusted p-value and log2FoldChange. Genesets from the DMPs were created by annotating the DMPs to their closest genes with *annotatePeak* from the ChIPseeker package [94]. STAT1 target score was calculated with the function *run_viper* from the dorothea package [105].

### Single-cell RNA-seq analysis and integration

For the single-cell RNA-seq analysis and integration with the bulk RNA-seq dataset we used the public dataset GSE174188 [76]. The dataset was converted to a Seurat object with the function *Convert* from SeuratDisk. Posterior analyses of the data were performed with Seurat package v4.3.0 [106]. Good quality cells were considered as those with a percentage of mitochondrial genes < 15%, a number of counts > 1500 and a number of features < 3500 and > 500. Analyses were performed only in the myeloid fraction of the total object. After filtering, myeloid cells were re-integrated (with *k.weight* of 50) taking into account batch information, doublets with T cells and platelets were removed. An object with 266k cells was obtained and used for clusterization and annotation. For Uniform Manifold Approximation and Projection (UMAP) representation, the functions *ScaleData*, *RunPCA*, *FindNeighbors*, *FindClusters* and *RunUMAP* were used sequentially, with the first 30 dimensions. The resolution used is indicated in each figure. Given the fact that the ethnicities represented in the bulk RNA-seq dataset were primarily European and Hispanic, for the bulk integration analyses (Figure 4D, H, I and Supplementary Figure 4A, B) these two cohorts were filtered resulting in a final object with 154k cells.

Clustering tree from the different resolutions was calculated and plotted with clustree package [107]. Top markers for each cluster was measured with the function findAllMarkers with the argument *min.cells.group* = 50. Gene Ontology of top 50 markers was calculated with *enrichGO* function from clusterProfiler package [103] using as background all the genes included in the analysis. Pseudotime was calculated with the package monocle3 [108–110]. Cell proportion from the bulk was measured with CIBERSORTx [78] and statistical inference of the proportions was measured with a linear model with the function *lm* and a two-sided Wilcoxon rank sum test.

## Data sharing

Methylation array and Expression data for this publication have been deposited in NCBI’s Gene Expression Omnibus and are accessible through GEO SuperSeries with accession number GSE249641.

## Acknowledgements

We thank CERCA Programme/Generalitat de Catalunya and the Josep Carreras Foundation for institutional support. The authors thank all the patients who graciously donated their time and samples for lupus research. E.B. was funded by the Spanish Ministry of Science and Innovation (MICINN; grant numbers PID2020-117212RB- I00 / AEI / 10.13038/501100011033). M.M.-G. was funded by Project PI18/00346 (Instituto de Salud Carlos III and co-funded by European Union (ERDF/ESF).

