## Supplementary Figure Legends for "Divergent Epigenetic and Transcriptomic Reprogramming of Monocyte Subpopulations in Systemic Lupus Erythematosus"

**Supplementary Figure 1.**  A. Summary violin plot representation of the last three clusters of DMPs. In grey, HD samples, in pink, SLE. B. Illustrative examples of two individual DMPs from the last three clusters of DMPs. In grey, HD samples, in pink, SLE. The color of the CpG name represents the cluster it belongs to. C and D. Barplot representation of DMPs annotation per cluster to CpG Island context (C) and gene body context (D). E. Motif enrichment analysis of the sequence englobing the DMPs from cluster M5, M6, M7. F. Gene Ontology enrichment results of the genes annotated the closest to the DMPs from the comparisons HD vs SLE in cMO and ncMO, respectively. Hypomethylated refers to lower methylation levels in SLE and hypermethylation to higher methylation levels in SLE. G. Gene Ontology enrichment results of the genes annotated the closest to the DMPs from the last three clusters of DMPs.

**Supplementary Figure 2**. A. Summary violin plot representation of the last three clusters of DEGs. In grey, HD samples, in pink, SLE. B. Illustrative examples of two DEGs from the comparison of HD vs SLE in each of the subsets. In grey, HD samples, in pink, SLE. C. Gene ontology enrichment of the last three clusters of DEGs. D. Expression of some highlighted transcription factors resulting from the motif enrichment analysis of DMPs. In grey, HD samples, in pink, SLE. E. Gene set enrichment analysis of the gene distribution between HD and SLE of genes annotating the closest to the DMPs in each subset. F. Summary of the gene set enrichment analysis statistical results.

**Supplementary Figure 3**. A and B. Violin plot representation of the iMO’s (A) and ncMO’s (B) CpGs correlating with SLE activity as measured by SLEDAI in a Spearman’s correlation. Samples divided in four groups according to their activity index: none is SLEDAI = 0, mild is SLEDAI between 1 and 5, moderate is an index between 6 and 10, and high is a SLEDAI above 10. C. Dotplot representation of methylation levels correlation with SLE activity index of the DMPs from clusters M1, M2, M3 and M4.

**Supplementary Figure 4**. A. Venn diagram representation of the overlap between the upregulated bulk DEGs (FDR < 0.05, log2FC > 1) and the single-cell DEGs (FDR < 0.05, log2FC > 0.1) in CD16- (left) and CD16+ (right) cells. B. Scatter plot representation of the correlation between the percentage of cells expressing each gene in the HD and SLE cohort in cluster 1 at the resolution of 0.05. In color the DEGs between SLE and HD samples in the single-cell dataset (FDR < 0.05, log2FC > 0.1). Labelled are the DEGs commonly upregulated in the bulk dataset comparison of HD versus SLE in either iMO or ncMO. C. Dotplot of representative gene ontology pathways enriched in the top 50 marker genes for each cluster generated at a resolution of 0.15.
